# A mono- and intralink filter (mi-filter) to improve false-discovery rates in cross-linking mass spectrometry data

**DOI:** 10.1101/2022.02.03.478943

**Authors:** Xingyu Chen, Carolin Sailer, Kai Michael Kammer, Julius Fürsch, Markus R. Eisele, Eri Sakata, Florian Stengel

## Abstract

Cross-Linking Mass Spectrometry (XL-MS) has become an indispensable tool for the emerging field of systems structural biology over the recent years. However, the confidence in individual protein-protein interactions (PPIs) depends on the correct assessment of individual inter protein cross-links. This can be challenging, in particularly in samples where relatively few PPIs are detected, as is often the case in complex samples containing low abundant proteins or in in-vivo settings. In this manuscript we are describing a novel mono- and intralink filter (mi-filter) that is applicable to any kind of crosslinking data and workflow. It stipulates that only proteins for which at least one monolink or intra-protein crosslink has been identified within a given dataset are considered for an inter-protein cross-link and therefore participate in a PPI. We show that this simple and intuitive filter has a dramatic effect on different types of crosslinking-data ranging from single protein complexes, over medium-complexity affinity enrichments to proteome-wide cell lysates and significantly lowers the number of false-positive identifications resulting in improved false-discovery rates for inter-protein links in all these types of XL-MS data.

## INTRODUCTION

An increasingly relevant approach for addressing protein-protein interactions (PPIs) is based on the rapidly evolving technology of crosslinking coupled to mass spectrometry (XL-MS). The general approach of protein XL-MS is based on covalent bonds that are formed using crosslinking reagents between proximal functional groups (most commonly lysine residues) in their native environment ^1-4^. The actual crosslinking sites are subsequently identified by mass spectrometry (MS) and reflect the spatial proximity of regions and domains within a given protein (intra-link) or between different proteins (inter-link). Additionally, the crosslinker can react twice within one peptide (loop-link) or only on one side with the peptide and hydrolyze on the other side (mono-link), revealing information on the accessibility of a specific amino acid residue. The field has seen significant technological and conceptual progress over the last couple of years and by now various enrichment strategies, different crosslinking chemistries and multiple detection and annotation strategies have been introduced^1, 2, 4^.

With the structural probing of recombinantly expressed static protein complexes by now being firmly established, it is the recent application of XL-MS on the systems level ^5^ and in living cells ^6^ that has spurred great interest and an ever-increasing number of studies ranging from bacterial, fungal and mammalian cell lysates and cultured cells^7, 8^, specific cellular organelles ^9-11^ and tissue^12, 13^ have been reported. These studies hint at the exciting prospect that XL-MS will soon be able to facilitate the structural probing of interaction partners of any protein of interest within living cells or even organisms.

However, the confidence in individual protein-protein interactions (PPIs) based on cross-linking data depends on the correct assessment of false discovery rates (FDRs) for individual inter protein cross-links. As recent data shows that FDRs in cross-linking data are frequently underestimated ^14^, which is particularly the case for inter-protein cross-links ^15^, this can undermine the confidence in individual PPIs and protein networks based on cross-linking data.

To date, FDR assessment in XL-MS has been primarily addressed through the optimization of scoring algorithms and the use of decoy databases ^11, 14, 16-18^. This approach works particularly well if large enough numbers of identifications can be reached for each particular cross-linking class (i.e. inter-protein, intra-protein and mon-link cross-links), but can be challenging in cases where relatively few PPIs are detected, as is often the case in complex samples containing low abundant proteins ^19^ or in in-vivo settings.

In this manuscript we therefore took a different approach and describe a novel mono- and intralink filter (mi-filter) that is applicable to any kind of crosslinking data and analysis pipeline. It stipulates that only proteins for which at least one monolink or intra-protein crosslink has been identified within a given dataset should be considered for an inter-protein cross-link and therefore participate in a PPI. We show that this simple and intuitive filter has a dramatic effect on all types of crosslinking-data ranging from single protein complexes, over medium-complexity affinity enrichments to proteome-wide settings and significantly reduces the number of false-positive identifications resulting in improved false-discovery rates in all these types of XL-MS data.

## RESULTS

### Concept of the Mi-filter

Our “mi-filter” (**M**onolink/**I**ntralink-filter) is based on the simple idea that only proteins for which at least one monolink or intra-protein crosslink has been identified within a given dataset should participate in an inter-protein cross-link and be part of a legitimate PPI (**Figure 1**). It is based on the observation that if the abundance of protein is high enough to be detectable by XL-MS, the formation rate of monolinks and intra-protein cross-links will be significantly higher than that of interlinks^19^. Or in other words, if no mono-link or intra-protein cross-link can be detected for a given protein, there is a high likelihood that this protein is not addressable by XL-MS in this particular sample and any inter-protein cross-link that includes this protein is likely a false-positive.

**Figure 1.**
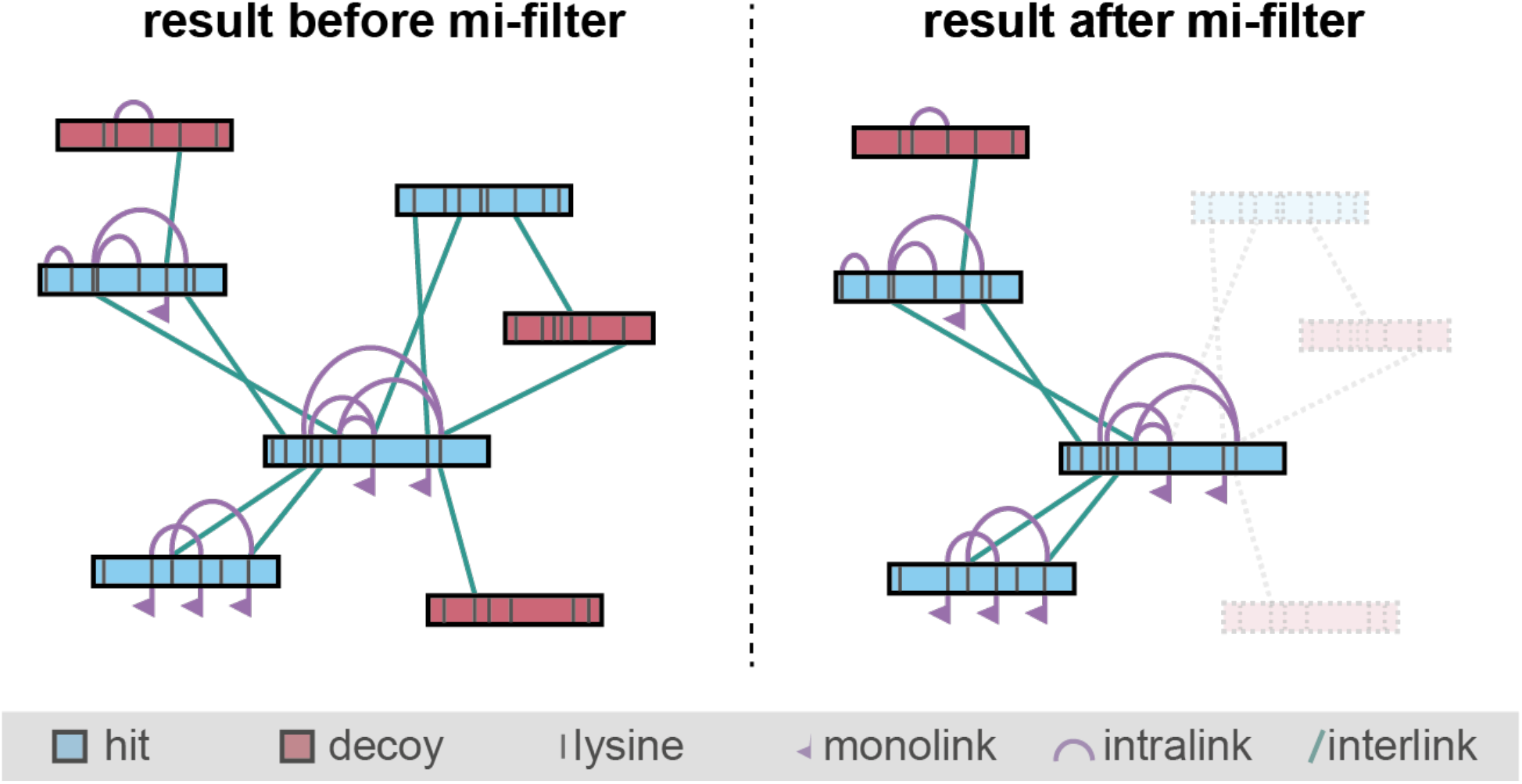
Concept of the mi-filter. Only proteins that contain at least one identified high-confidence monolink or intra-protein crosslink are considered for inter-proteins cross-links and can therefore be part of a PPI.

Our approach is therefore not designed as a contradiction to prevalent FDR calculations based on scoring algorithms and the use of decoy databases but is rather intended as an additional tool that can be applied on top of existing workflows in order to minimize false-positive assignments of inter-protein cross-links or as an alternative, in case no sufficient data coverage for conventional FDR calculations can be achieved.

### Inter-protein cross-links are disproportionally affected by high FDRs

Minimizing false-positive assignments for inter-protein cross-links is particularly crucial as all PPIs based on cross-linking data depend entirely on information from inter-protein crosslinks. Moreover, inter-protein cross-links are disproportionally affected from high FDRs (**Figure 2**). **Figure 2** shows the amount of detected decoy hits for monolinks, intra-protein and inter-protein cross-links for the 26S proteasome at increasingly stringent filtering settings, i.e. increasing agreement between measured experimental and in-silico generated reference spectra. The data shows that the relative proportion of detected decoy hits for inter-protein cross-links is significantly larger than for mono or intra-protein links for all settings and, importantly, that inter-protein cross-links still contain a significant number of false-positive identifications at cut-offs where the number of detected decoys (and thus resulting FDRs) for intra-protein and monlinks are already negligible.

**Figure 2.**
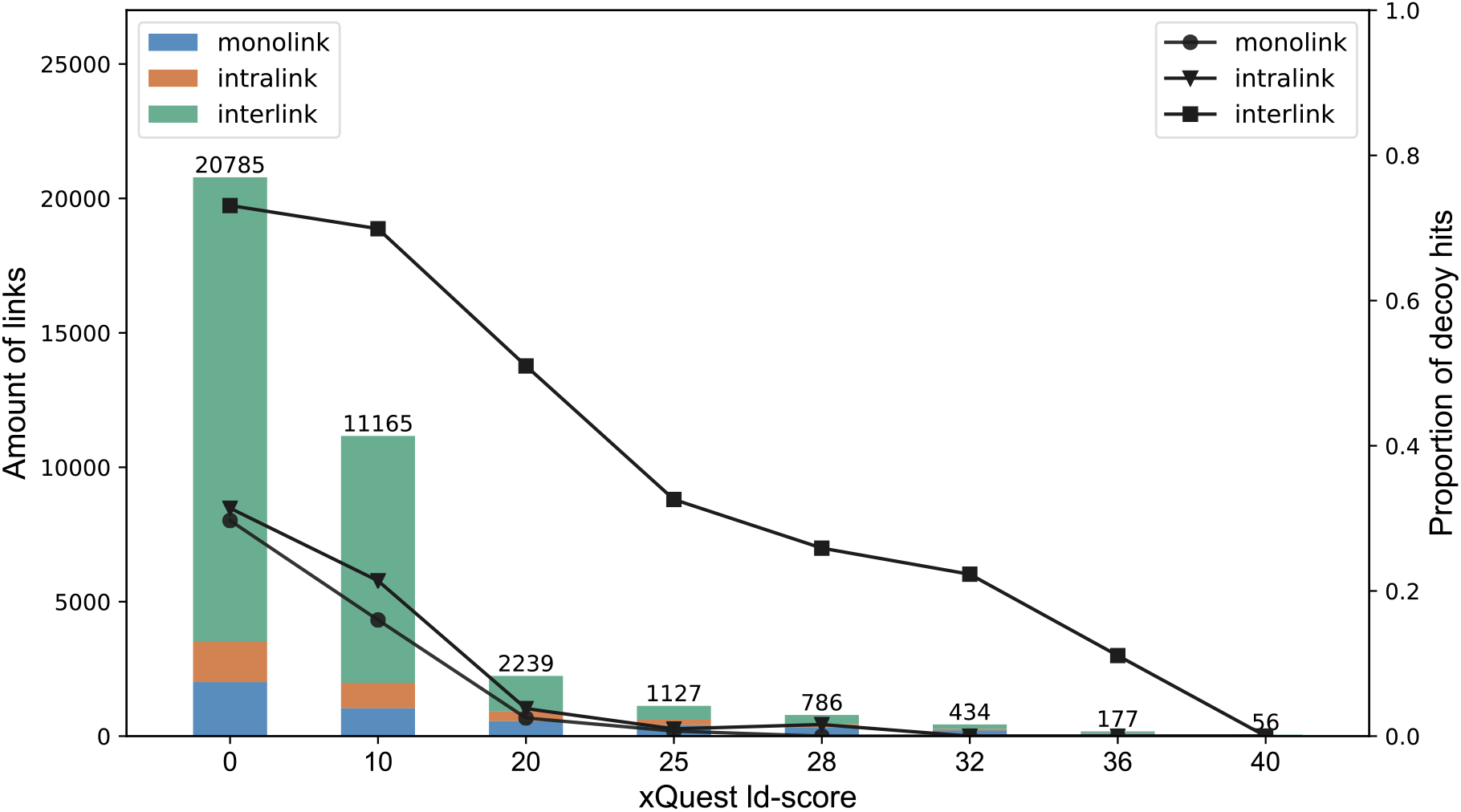
Inter-protein cross-links are disproportionally affected from high FDRs. The bar chart shows the cumulative amount of detected monolinks, intra-protein cross-links and inter-protein cross-links (y axis, left) of the 26S proteasome at different ld-score cut-offs ^22^, i.e. increasing levels of agreement between measured experimental and in-silico generated reference spectra. Blotted are the relative proportion of detected decoy hits (y-axis, right) versus the respective ld-score setting (x-axis).

Therefore, in cases where FDRs are assigned in toto over all classes of cross-links or in cases where FDRs based on decoy-measurements cannot be properly calculated, for example due to sparsity of the data, the likelihood of a false-positive assignment for inter-protein cross-links increases dramatically.

### *Mi-filter* improves inter-protein FDRs for different types of cross-linking data

In order to evaluate the effect of the *mi-filter* on FDRs, we applied it to typical cross-linking datasets of different complexity using the *xQuest/XProphet* ^20^ pipeline as an example (**Figure 3**). Our least complex sample is the 26S proteasome from *S. cerevisiae* consisting of 34 proteins (**Figure 3A** and **B**). An intermediate one is the combined dataset of pre-60S ribosomal particles obtained by affinity enrichment, containing a total of around 300 proteins (**Figure 3C** and **D**). We could previously show that the application of the mi-filter to this dataset results in significantly improved FDRs ^21^, but only now we have thoroughly investigated the influence of the *mi-filter* on this and other datasets and for various settings of increasingly stringent filtering. The most complex sample we evaluated using our *mi-filter* is a proteome-wide XL-MS dataset of *S. cerevisiae* cell lysate^19^ (**Figure 3E** and **F**).

**Figure 3.**
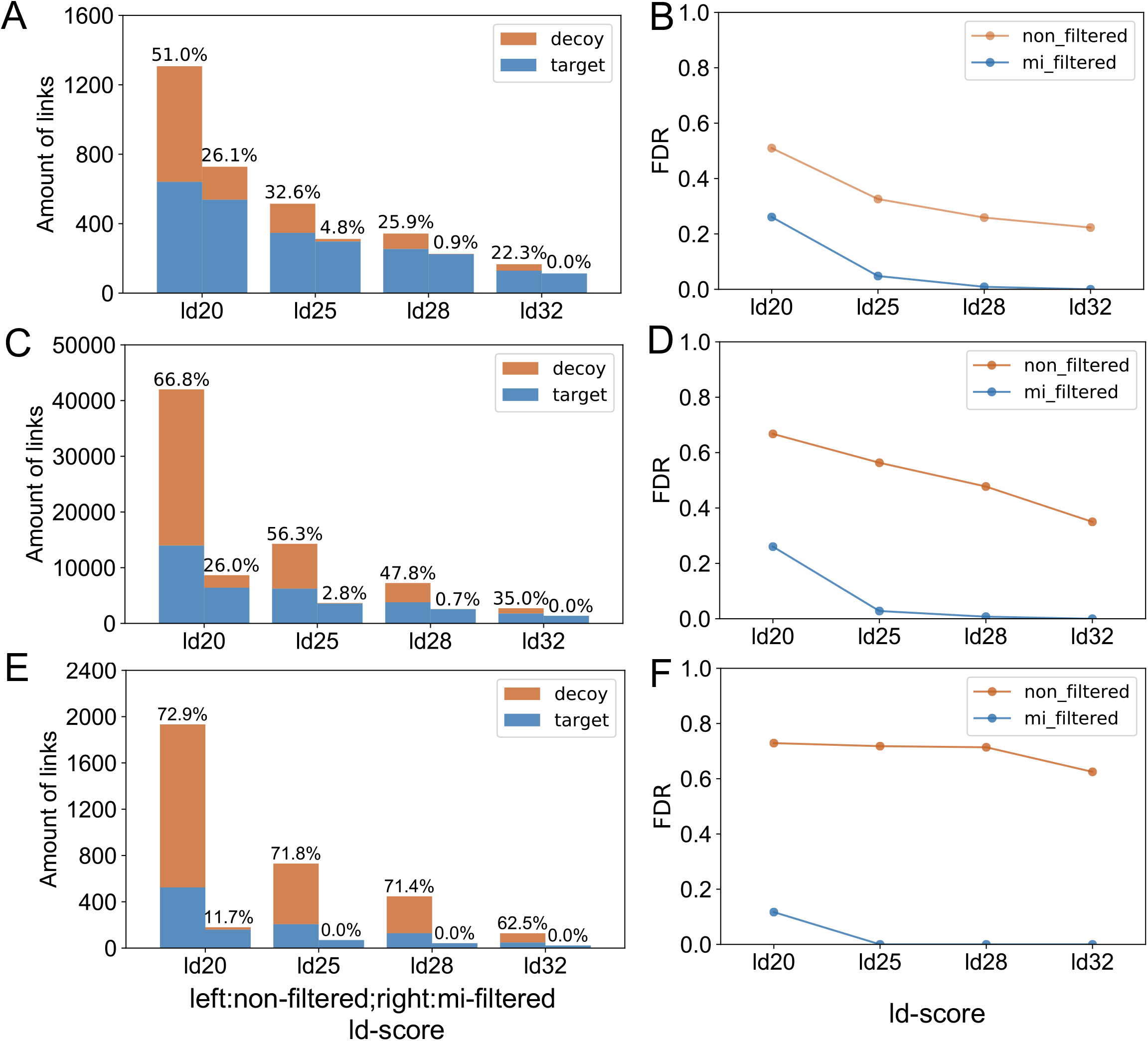
Comparison of FDRs with and without mi-filter. Inter-protein cross-links are shown for three different types of datasets representing typical experimental set-ups including the 26S proteasome from *S. cerevisiae* as an example of a single protein complex (**A** and **B**), affinity enrichments of pre-60S ribosomal particles (**C** and **D**) and a proteome-wide cross-linking experiment using *S. cerevisiae* cell lysate (**E** and **F**). Panels **A, C** and **E** show the number of decoy and target hits (non-decoy hits) in non-filtered (left bar in each group) and *mi-filtered* samples (right bar in each group) for increasingly stringently filtered data. Target hits are shown in blue and decoys in red. The relative percentage of decoy hits for each setting is indicated. Panels **B, D** and **F** show the false discovery rate (FDR) of non-filtered (red line) versus mi-filtered results (blue line) for the respective datasets. The FDR was calculated as the proportion of decoy hits to the total amount of all hits (target plus decoy hits).

We first had a closer look at the relative abundance of proteins that were filtered-out by the *mi-filter*, taking the pre-60S ribosomal particles dataset as an example ^21^. Here proteins for which a mono- or intra-protein link was detected are in average of significantly higher abundance than proteins without mono- or intra-protein links (**Supplementary Figure 1**). This already indicates that proteins without mono- or intra-protein links are either not present in the sample at a concentration high enough for crosslink identification or they are not present at all.

After application of the *mi-filter* (right bar of each group) all datasets consistently exhibit a significant decrease in the number of detected decoy inter-protein links (**Figure 3A, 3C** and **3E**) as well as a significant decrease in the resulting FDRs (**Figure 3B, 3D** and **3D**). It is interesting to note that this is not only true for the different sample types but also for the increasingly stringent filtering settings (i.e. increasingly good matches between experimental data and and in-silico generated reference spectra). The *mi-filter* is therefore not only able to filter out most decoy links already at medium filtering settings (**Figure 3B, 3D** and **3D**), it also is able to retain the majority of bona-fide true positive links, as indicated by the diminishing difference of detected target hits between filtered and non-filtered data for the highest quality MS data (**Figure 3A, 3C** and **3E**).

Taken together this demonstrates the value of the *mi-filter* as a stringent filtering device that results in a significant reduction of decoy inter-protein cross-link identifications and improved FDRs for inter-protein cross-links in different types of cross-linking data.

### Quality of the mi-filtered data

In a next step we wanted to test and benchmark the *mi-filter* also for its ability to identify true-positive cross-links in a proteome-wide setting. In contrary to mixtures of purified proteins or protein complexes, which can be benchmarked against existing atomistic high-resolution structures in order to obtain true-positive identifications (which differs from the mere calculation of FDRs from decoy-hits), there is no known ground-truth in a proteome-wide cross-linking experiment, as the precise protein arrangement within a cell or lysate is unknown.

We therefore took the totality of MS and MS/MS spectra that we had experimentally obtained from a sample of crosslinked 26S proteasome and searched it in a proteome-wide setting (i.e. against a large protein database; *see* methods for details) with and without application of the *mi-filter* (**Figure 4A**). Our data shows that the application of the *mi-filter* did not only lead to a significant reduction of detected decoy-hits and a nearly 35-fold reduction in the resulting FDR. When mapped onto the published high-resolution cryo-EM structure of the *S. cerevisiae* 26S proteasome (PDB 4CR2), over 90 % of our mi-filtered cross-links fall within 35 A°, the maximal lysine Ca-Ca distance that our crosslinker can bridge (**Supplementary Figure 2**), indicating that our *mi-filtered* cross-links are also biologically meaningful.

**Figure 4.**
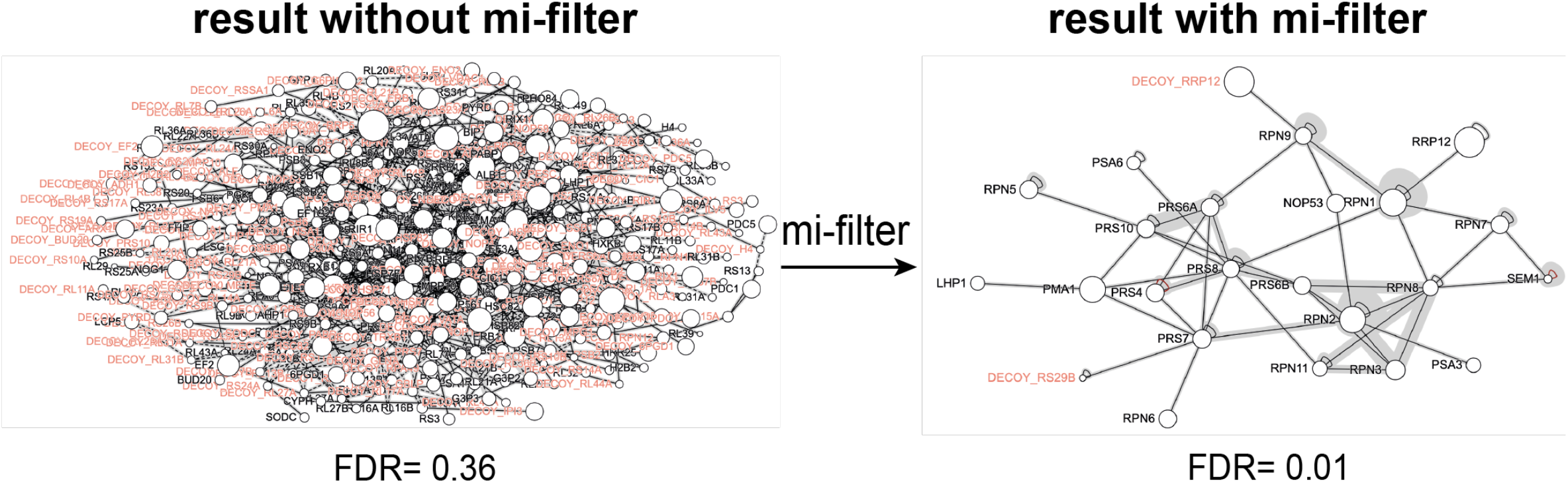
Quality of the mi-filtered data. Shown is a cross-linking dataset of the 26S proteasome with and without application of the *mi-filter* searched against a manually curated database mimicking proteome-wide protein distributions (*see* methods for details).

In summary our data indicates that the *mi-filter* is not only able to significantly reduce the amount of detected decoy-hits and resulting FDRs but is simultaneously able to identify and retain true-positive cross-links also in a proteome-wide setting.

## DISCUSSION

In this manuscript we describe and benchmark a novel mono- and intralink filter that is in principle applicable to any kind of crosslinking data and analysis pipeline. This simple and intuitive *mi-filter*, which removes inter-protein cross-links if the connected polypeptides are not additionally represented within their respective intralink or monolink pools, reduces identification of false-positives and improves FDRs for inter-protein cross-links significantly. We show that this is true for different types of crosslinking-data ranging from single protein complexes, over medium-complexity affinity enrichments to proteome-wide settings. Moreover, in addition to reliably reducing the amount of detected decoy-hits in a given cross-linking sample, the *mi-filter* is also able to identify and retain the majority of true-positive cross-links, also in a complex proteome-wide setting.

While we have used the *xQuest/xProphet* cross-linking software suite in this manuscript as an example to benchmark the *mi-filter*, our *mi-filter* can in principle be applied to any cross-linking software to further improve and validate the obtained cross-linking data. It can also be used as a stand-alone tool to minimize false-positive assignments and improve the detection of true and biologically relevant PPIs. This is particularly useful in cases where no sufficient data for conventional FDR calculations based on decoys can be achieved, as is often the case in complex samples containing low abundant proteins or in *in-vivo* settings.

Taken together our *mi-filter* greatly enhances the reliability of individual inter-protein cross-links in any type of cross-linking data and therefore their ability to provide reliable and biologically relevant positional information as source of a PPI.

## Acknowledgements

This work was supported by the German Research Foundation through Germany’s Excellence Strategy - EXC 2067/1-390729940, SFB1035/Project A01, and CRC889/Project A11 to E.S. F.S. is grateful for funding from an ASAP Collaborative Research Network Grant (ASAP-000519) and the German Research Foundation (STE 2517/1 and STE 2517/5-1).

## Author Contributions

X.C., C.S. and F.S. conceived the study and experimental approach. M.R.E and E.S. expressed and purified the 26 S proteasome. X.C. and C.S. implemented and applied the mi-filter with help from K.M.K. J.F. performed original proteome-wide studies. C.S. carried out affinity-enrichments. X.C., C.S., K.M.K and F.S. analysed the data. X.C., C.S. and F.S. wrote the paper with input from all authors.

## Materials & Methods

### Mi-filter script

The *mi-filter* script was written in python and is available at the Github repository (https://github.com/xingyu-konstanz/mi-filter.git). It is tailored to *xQuest*^20^ output tables but can in principle be applied to crosslinking MS datasets obtained by any of the established crosslink-identification software platforms such as MeroX^23^, Xlinkx^17^, Xi^24^, pLink2^25, 26^or RNPxl^27^. It selects proteins from the input dataset, which contain at least one mono- or intra-protein link and subsequently filters for inter-protein crosslinks within this list. It also calculates a simple FDR (using the ratio of target and decoy links) at each ld-Score cut-off for monolinks, inter-protein and intra-protein cross-links separately. In detail, the *mi-filter* script works as follows: it filters the input files for a specified ld-Score, it then concatenates the input data frames and, if specified, filters for biological replicates of cross-linking sites (therefore input files must be sorted by biological replicates). In a next step it adds a ‘decoy’ column to the concatenated data frame. In the ‘XLtype’ column, strings are replaced in a way that only three types of cross-link are left: monolinks, intra-protein and inter-protein cross-links. Proteins without a monolink or an intra-protein cross-link are then filtered out by the parameter “–mi “when running the mi-filter program. Subsequently, a simple FDR

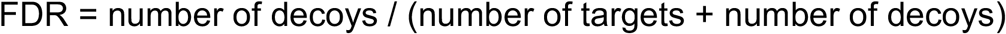

is calculated at each ld-Score cut-off (using 1.0 score units) for monolinks, intra-protein and inter-protein cross-links separately. Please note that links containing at least 1 decoy peptide are considered as decoys and there is no discrimination between cross-links of the following type T-T, T-D, D-T and D-D (where T denotes a target peptide and D a decoy peptide).

### 26S proteasome cross-linking dataset

Purification of yeast 26S proteasomes was performed as described in ^28^. *S. cerevisiae* cells (YYS40; MATa rpn11::RPN113FLAG-HIS3) were grown for 48 hours and harvested in stationary phase. The purification of 3XFLAG-tagged 26S proteasome was carried out by affinity purification using M2 anti-FLAG beads (Sigma A2220). After incubation for 1.5 hrs at 4°C the proteasome was eluted with FLAG peptide. An overnight sucrose gradient was carried out for the second purification step. The sucrose gradient was centrifuged in a Beckman SW41 rotor for 17 h at 4°C at 28000 rpm. Proteasome-containing fractions were identified by degradation of the peptide suc-LLVY-AMC, SDS-PAGE analysis, and Bradford assay.

100 µg of purified 26S proteasome (1µg/µl) were subsequently incubated with the isotopically labeled crosslinking reagent disuccinimidyl suberate d0/d12 (DSS-H12/D12, *Creativemolecules* Inc.) at a final concentration of 1 mM for 30 min at 30 °C while shaking at 650 rpm in a Thermomixer (Eppendorf). The reaction was quenched with ammonium bicarbonate at a final concentration of 50 mM for 10 min at 30 °C and 650 rpm. Crosslinked samples were dried (Eppendorf, Concentrator plus), resuspended in 100 µl 8M Urea, reduced, alkylated, and digested with trypsin (Promega). Digested peptides were separated from the solution and retained by a solid phase extraction system (SepPak, Waters). Crosslinked peptides were enriched by size exclusion chromatography using an ÄKTAmicro chromatography system (GE Healthcare) equipped with a Superdex™ Peptide 3.2/30 column (column volume = 2.4 ml). Fractions were collected in 100 µl units and analyzed by LC-MS/MS. For each crosslinked sample two fractions (1.2-1.3 ml and 1.3-1.4 ml) were collected and measured in technical duplicates. Absorption levels at 215 nm of each fraction were used to normalize peptide amounts prior to LC-MS/MS analysis.

LC-MS/MS analysis was carried out on an Orbitrap Fusion Tribrid mass spectrometer (Thermo Electron, San Jose, CA). Peptides were separated on an EASY-nLC 1200 system (Thermo Scientific) at a flow rate of 300 nl/min over an 80 min gradient (5 % acetonitrile in 0.1 % formic acid for 4 min, 5 % - 35 % acetonitrile in 0.1% formic acid in 75 min, 35 % - 80 % acetonitrile in 1 min). Full scan mass spectra were acquired in the Orbitrap at a resolution of 120,000, a scan range of 400 - 1500 m/z, and a maximum injection time of 50 ms. Most intense precursor ions (intensity ≥ 5.0e3) with charge states 3 - 8 and monoisotopic peak determination set to ‘peptide’ were selected for MS/MS fragmentation by CID at 35 % collision energy in a data dependent mode. The duration for dynamic exclusion was set to 60 s. MS/MS spectra were analyzed in the iontrap at a rapid scan rate.

For the crosslink identification of the 26 proteasome in a “proteome-wide setting” a database was compiled which contained the 34 proteins of the 26S proteasome plus the 200 most abundant proteins in *S. cerevisiae* as annotated in the PAX database (https://pax-db.org/). MS raw files were subsequently converted to centroid files and searched using *xQuest* in ion-tag mode. Crosslinks were exported as .tsv files with the filter settings deltaS = 95 and a max. ppm range from -5 to 5, containing all (non-unique) identifications. The *mi-filte*r was applied to different ld-Score cut-offs (20, 25, 28 and 32) and FDRs were calculated as described above and compared to the dataset before mi-filtering (**Supplementary Data 1** and **Figure 3A** and **3B**).

### Pre-60S ribosome XL-MS dataset

The dataset consists of biological triplicate measurements of 12 different pre-60S ribosomal particles, which were enriched using affinity-tagged RBFs and was collected as part of another study ^21^. The *mi-filte*r was applied to different ld-Score cut-offs (20, 25, 28 and 32) and FDRs were calculated as described above and compared to the dataset before mi-filtering (**Supplementary Data 2** and **Figure 3C** and **3D**).

### Proteome wide cross-linking dataset

The dataset contains biological triplicate measurements of cell lysate in *Saccharomyces cerevisiae* and was collected as part of another study^19^. *xQuest* results from this paper were directly downloaded and the cross-linked sample using equimolar concentrations (1x) of BS3 as a cross-linker was chosen for further analysis. The *mi-filter* was applied to different ld-Score cut-offs (20, 25, 28 and 32) and FDRs were calculated as described above and compared to the dataset before mi-filtering (**Supplementary Data 3** and **Figure 3E** and **3F**).

### Mapping of filtered crosslinks

Crosslink networks were visualized with xiNET ^29^.

## Data availability

The MS raw files, the crosslink databases and original *xQuest* result files have all been deposited to the ProteomeXchange Consortium via the PRIDE partner repository ^30^ with the project accession number PXD031215 (Username: Password: yLjErDmt). The previously published ribosome ^21^ and lysate ^19^ datasets have the project accession numbers PXD021831 and PXD014759, respectively.

## Supplemental Figures

**Supplementary Figure 1:**
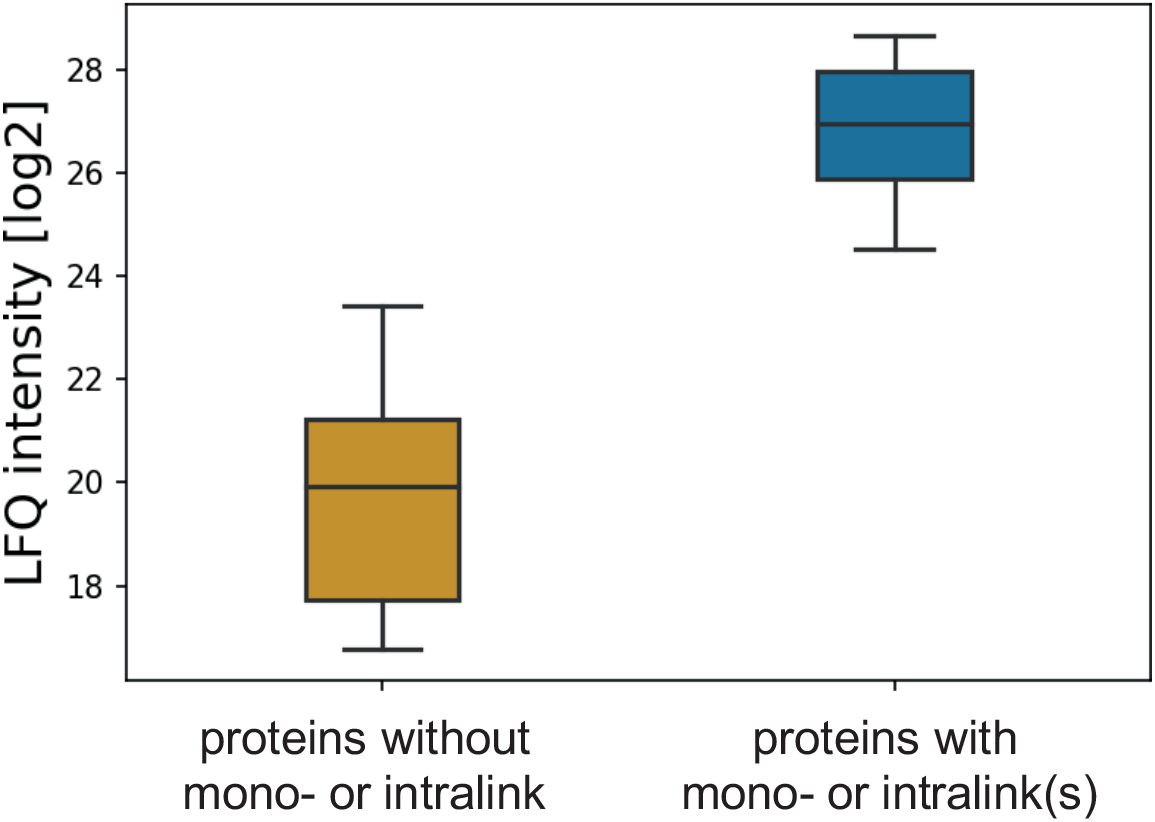
Relative Abundance of proteins without mono- or intra-protein link that were filtered-out by the *mi-filter* (yellow) compared to the relative abundance of proteins with a mono- or intra-protein link (blue). LFQ intensities were calculated based on peptides which were not modified by the crosslinking reagent DSS.

**Supplementary Figure 2:**
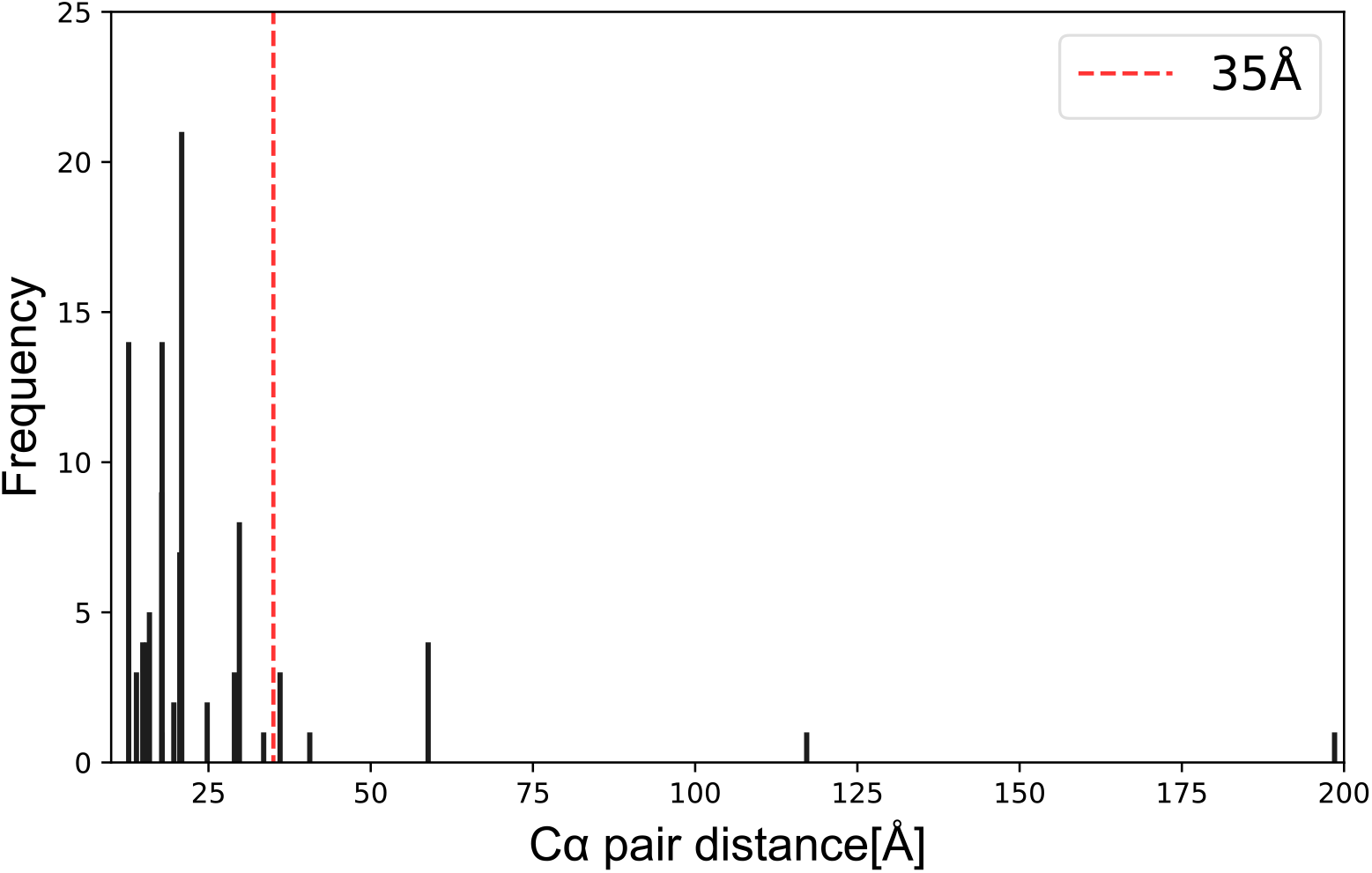
Histogram of mapped distances of all cross-linked residues within the mi-filtered 26S proteasome data set versus their frequency. The 35A° threshold is indicated as a red dotted line.

